# Variants with the N501Y mutation extend SARS-CoV-2 host range to mice, with contact transmission

**DOI:** 10.1101/2021.03.18.436013

**Authors:** Xavier Montagutelli, Matthieu Prot, Laurine Levillayer, Eduard Baquero Salazar, Grégory Jouvion, Laurine Conquet, Maxime Beretta, Flora Donati, Mélanie Albert, Fabiana Gambaro, Sylvie Behillil, Vincent Enouf, Dominique Rousset, Hugo Mouquet, Jean Jaubert, Felix Rey, Sylvie van der Werf, Etienne Simon-Loriere

**Affiliations:** Institut Pasteur, Université de Paris, Mouse Genetics Laboratory, F-75015 Paris, France; Institut Pasteur, Université de Paris, G5 Evolutionary Genomics of RNA Viruses, F-75015 Paris, France; Institut Pasteur, Université de Paris, Functional Genetics of Infectious Diseases Unit, F-75015 Paris, France; Institut Pasteur, Université de Paris, Structural Virology Unit, F-75015 Paris, France; Ecole nationale vétérinaire d’Alfort, Unité d’Histologie et d’Anatomie Pathologique, Maisons-Alfort, France; Université Paris Est Créteil, EnvA, ANSES, Unité DYNAMIC, Créteil, France; Institut Pasteur, Université de Paris, Humoral Immunology Laboratory, F-75015 Paris, France; Institut Pasteur, Université de Paris, Molecular Genetics of RNA viruses Unit, CNRS UMR 3569, F-75015 Paris, France; Institut Pasteur, Université de Paris, National Reference Center for Respiratory Viruses, F-75015 Paris, France; Institut Pasteur de la Guyane, Laboratoire de Virologie, Cayenne, French Guiana, France

**Keywords:** SARS-CoV-2, variants, host range, mice, transmission, reservoir

## Abstract

Receptor recognition is a major determinant of viral host range, infectivity and pathogenesis. Emergences have been associated with serendipitous events of adaptation upon encounters with novel hosts, and the high mutation rate of RNA viruses may explain their frequent host shifts. SARS-CoV-2 extensive circulation in humans results in the emergence of variants, including variants of concern (VOCs) with diverse mutations notably in the spike, and increased transmissibility or immune escape. Here we show that, unlike the initial and Delta variants, the three VOCs bearing the N501Y mutation can infect common laboratory mice. Contact transmission occurred from infected to naive mice through two passages. This host range expansion likely results from an increased binding of the spike to the mouse ACE2. Together with the observed contact transmission, it raises the possibility of wild rodent secondary reservoirs enabling the emergence of new variants.

## INTRODUCTION

Host range expansion or switch to other species has been prevalent in the course of coronaviruses evolutionary history (Cui et al., 2019; Woolhouse et al., 2005). Understanding the host range and how it is modified as the pathogen evolves is critical to estimate the emergence risk and determine the reservoirs to monitor. In the case of the severe acute respiratory syndrome coronavirus 2 (SARS-CoV-2), responsible for the ongoing coronavirus disease 2019 (COVID-19) pandemic, animals such as non-human primates, hamsters, ferrets, minks and cats were shown to be permissive (Johansen et al., 2020). By contrast, the zoonotic virus was shown to not replicate in mice and rats due to poor binding of the virus spike on the rodent cellular receptor angiotensin-converting enzyme 2 (ACE2) (Zhou et al., 2020).

At the end of 2020, the emergence of variants of concern (VOCs) was noted in different parts of the world (WHO, 2021). The Alpha variant (Pango lineage B.1.1.7 (Rambaut et al., 2020a)), was noted for its rapid spread in the UK (Rambaut et al., 2020b), while the Beta variant (Pango lineage B.1.351) expanded in multiple regions of South Africa (Tegally et al., 2021), and the Gamma variant (Pango lineage P.1) emerged in Manaus, Brazil (Faria et al., 2021). The global circulation and spread of these variants have led to concerns about increased transmission and their potential to evade immunity elicited by vaccination or naturally acquired. More recently, the Delta variant (Pango lineages B.1.617.x and AY.x), initially described in India, has rapidly expanded worldwide and became dominant in many countries (Mlcochova et al., 2021).

Alpha, Beta and Gamma SARS-CoV-2 VOCs all harbor the N501Y change in the spike glycoprotein which belongs to a set of 6 key amino acid residues critical for the tight interaction of the receptor binding domain (RBD) with hACE2 (Yi et al., 2020). Strikingly, this mutation was also noted, among others, in independently generated mouse-adapted SARS-CoV-2 strains (Gu et al., 2020; Rathnasinghe et al., 2021). Here, we assessed the replication potential in cells and in mice of low-passage clinical SARS-CoV-2 VOCs isolates as well as their transmission potential by direct contact between mice.

## RESULTS

### The SARS-CoV-2 N501Y variants efficiently replicate in mice airways

We first measured the capacity of a panel of low-passage clinical isolates belonging to the initial lineages and to each of the VOCs lineages (Fig. S1) to infect mouse cells expressing the murine ACE2 receptor (DBT-mACE2), by comparison with VeroE6 cells. Unlike the ancestral viruses (B and B.1 lineages) which did not replicate in these cells, all VOCs carrying the N501Y change replicated to high titers at 48h post infection (Fig. S2).

We next inoculated 8-week-old BALB/c and C57BL/6 mice intranasally (i.n.) with a representative virus of either the most prevalent SARS-CoV-2 lineage in 2020 (basal to B.1, carrying the D614G substitution) or the four VOCs lineages and we measured the viral load and titer in the lungs on day 3 post infection (dpi3). Consistently with what was reported for the ancestral virus (Dinnon et al., 2020), low amounts of viral RNA and no infectious particles were detected with the D614G (B.1) virus (Fig. 1A-B). Similar observations were made with a Delta variant. In contrast, inoculation with Beta or Gamma variants yielded virus replication to high titers in lung tissues in both mouse strains. A strain of the A.27 lineage, harboring the N501Y change without the D614G mutation, similarly yielded comparably high viral titer in the lungs of C57BL/6 mice. Interestingly, while the Alpha strain resulted in significantly lower viral load than Beta or Gamma, an Alpha variant carrying the E484K mutation in the spike (present in Beta and Gamma) yielded viral titer comparable to the other N501Y isolates.

**Fig. 1.**
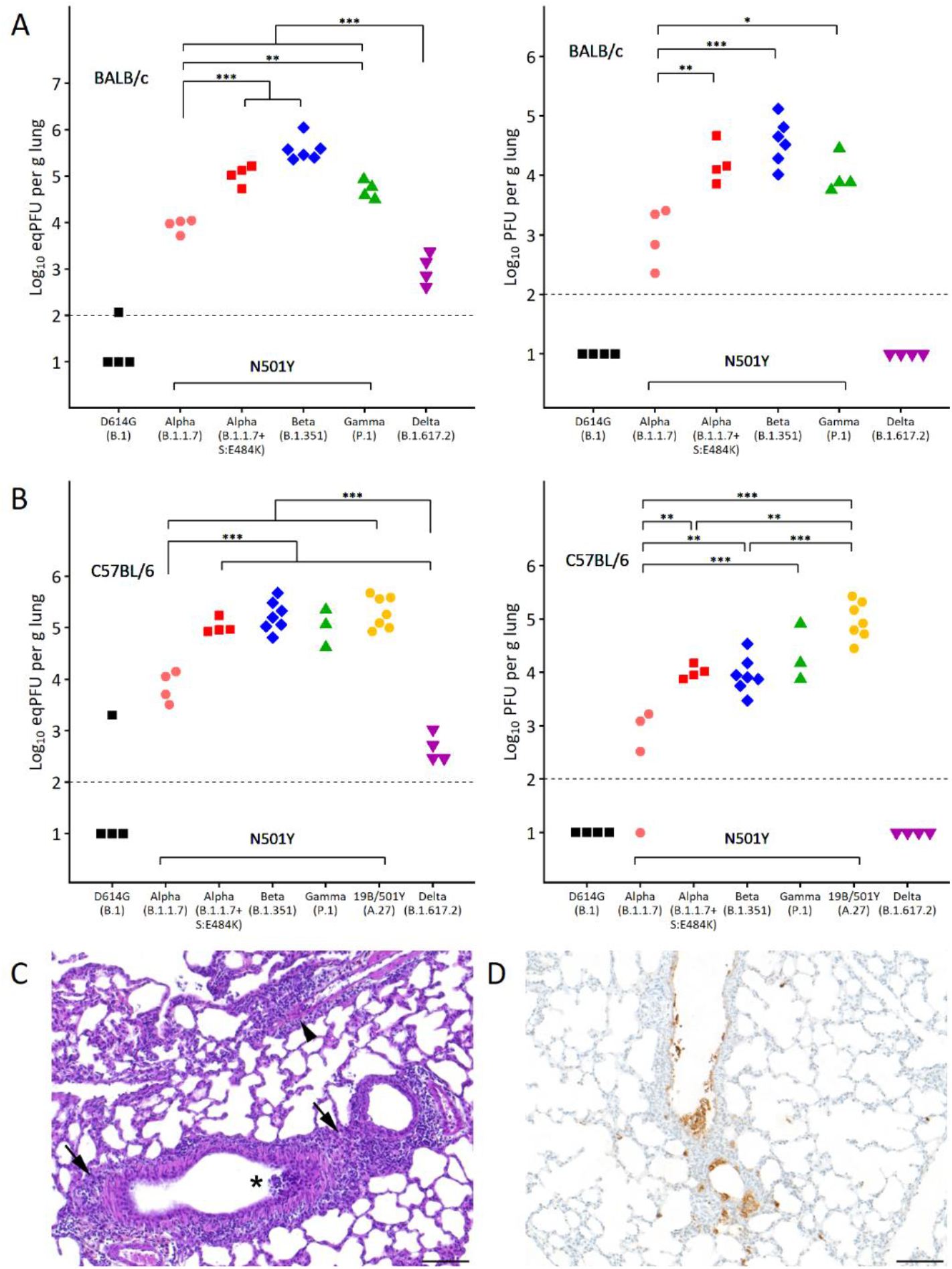
Variants of SARS-CoV-2 carrying the N501Y mutation replicate to high titers in the lungs of young adult mice. Eight-week-old BALB/c (A) and C57BL/6 (B) mice were inoculated intranasally with 6×10^4^ PFU of SARS-CoV-2 isolates. Lungs were harvested at 3 dpi. Viral load was quantified by RT-qPCR. Viral titer was quantified on VeroE6 cells. The dotted line represents the limit of detection (LOD), and undetected samples are plotted at half the LOD. (C-D) Lung of a BALB/c mouse inoculated with the Beta variant. (C) High magnification of the lung histological lesions: multifocal infiltrates of inflammatory cells (mononuclear cells and neutrophils) in the interstitium around bronchi/bronchioles (arrows) and blood vessels (black arrowhead), with focal destruction of bronchiolar epithelial cells (*), HE staining. (D) Detection of multifocal infected cells by anti N-immunohistochemistry (scale bar: 100 μm).

To capture the time course of the productive infection with the Beta and Gamma viruses, we also sampled infected mice at day 2 and 4, revealing an early peak of infection (Fig. S3). None of these variants induced clinical signs and only the Beta variant induced a moderate and transient body weight loss at dpi2-3 (Fig. S4).

Three days after Beta or Gamma virus inoculation, histological evaluation of the lung ranged from normal morphology to moderate lesions characterized by multifocal interstitial infiltrates of lymphocytes, plasma cells, macrophages and rare neutrophils (around bronchi/bronchioles and blood vessels) and by degenerating epithelial cells in the bronchial and bronchiolar spaces (Figure 1C). Anti-N immunohistochemistry revealed the presence of infected cells in bronchiolar epithelium, bronchiolar and alveolar spaces and alveolar walls (Figure 1D).

### The Beta variant can disseminate in mice by close contact

The extension of SARS-CoV-2 host range to mice raises the possibility that wild rodents become secondary reservoirs. To assess the capacity of N501Y-bearing variants to disseminate in mouse populations, we inoculated mice with the Beta variant and introduced naive mice (named Contact 1) in their cage one day post-infection in 3 independent replicates (Figure 2A). Two or three days later, we removed the naive mice and placed them in a cage with other naive mice (Contact 2) to assess secondary transmission. On day 17, infected mice had high titers of anti-S as revealed by ELISA, and Contact 1 and Contact 2 mice had similar intermediate levels (Figure 2B), indicating that both contact groups had been exposed to the virus.

**Fig. 2.**
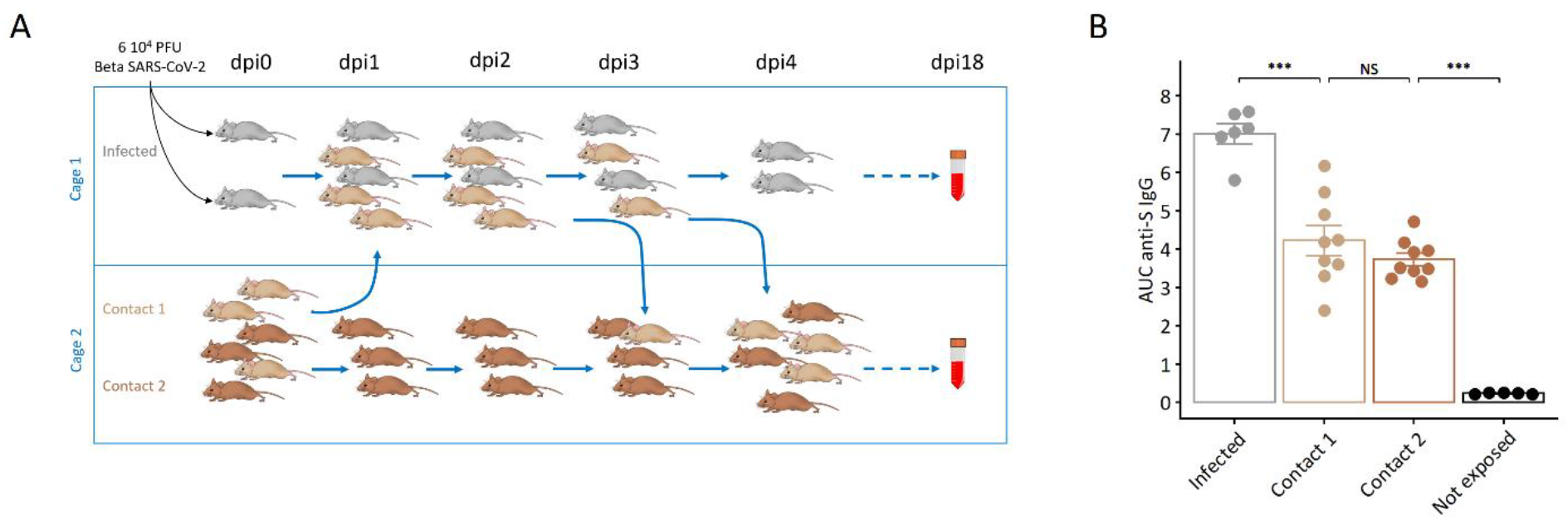
Transmission of SARS-CoV-2 Beta variant in mice by direct contact. (A) Two eight-week-old BALB/c mice were inoculated intranasally with 6×10^4^ PFU of Beta SARS-CoV-2. On the following day (dpi1), three naive Contact 1 mice were introduced in their cage. One Contact 1 mouse was removed on dpi3 and placed in a cage containing three naive Contact 2 mice. On dpi4, the two remaining Contact 1 mice were moved to cage 2 with the other Contact 1 and the Contact 2 mice. Two weeks later (dpi18), anti-S antibody level was assessed by ELISA in all mice. The experiment included three such cohorts. (B) ELISA results as area under the curve (AUC). Infected mice (total n=6) had the highest antibody titers (7.0 ± 0.26; mean ± sem). Contact 1 and Contact 2 mice showed similar levels of antibodies (4.22 ± 0.38 and 3.73 ± 0.17, respectively; p > 0.54 ANOVA with Tukey’s multiple comparisons test) while unexposed mice had background levels (0.23 ± 0.01), indicating that all mice of both contact groups had been exposed to the virus.

**Fig. 3.**
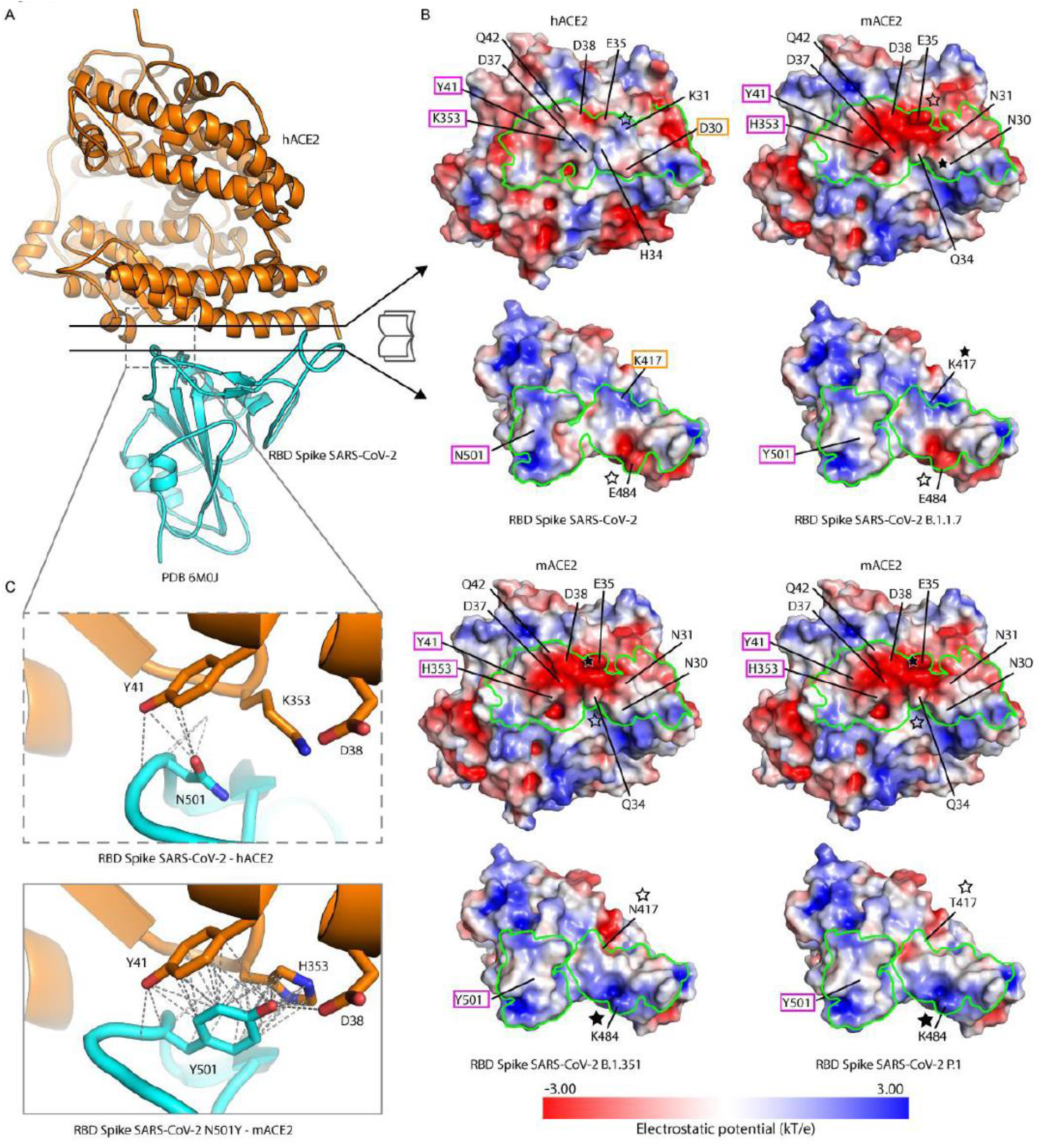
Homology modeling of mouse ACE2 in complex with the RBD of SARS-CoV-2 variants. A. Ribbon representation of the overall structure of the peptidase domain of hACE2 (orange) bound to SARS-CoV-2 spike RBD (cyan). **B.** Open book representations of the interacting surfaces between hACE2 and SARS-CoV-2 spike RBD (crystal structure 6M0J) and mACE2 with the RBDs of the B.1.1.7, B.1.351 and P.1 variants (homology models). The surfaces are colored by electrostatic potential from red (negative charge, −3.00 kT/*e*) to blue (positive charge, 3.00 kT/*e*), as indicated by the colored bar underneath. The green outline indicates the contact area between the two molecules. The residues of the central negatively charged patch in ACE2 and the mutation sites in the RBDs are labelled. The residues in the RBD and in ACE2 that come into contact in the complex are labelled within boxes of the same color. The stars indicate the RBD residues and their vicinity area in the interaction with ACE2. **C.** Detailed views of the interaction of the RBD residue N501 with hACE2 (PDB 6M0J, upper inset) and the mutant Y501 with mACE2 (homology model, lower inset). The dashed lines indicate possible electrostatic interactions between side and main chains. The model of the interaction of the N501Y RBD mutant with mACE2 indicates that the tyrosine side chain lies between the mACE2 residues Y41 and H353, making π-π interactions and a hydrogen bond with D38, suggesting a stronger interaction between mACE2 and RBDs carrying this mutation.

### Structural changes in the spike affect viral replication in mice

Homology modeling of the spike in contact with mACE2 reveals that the murine version of the receptor presents a strongly negatively charged central patch composed of residues E35, D37, D38 and Q42. These residues are also present in hACE2 but associated with several positively charged residues, which are replaced by more neutral amino acids in mACE2 (Figure 2B). This feature indicates that efficient binding to mACE2 would require complementary positive charges in the RBD. As expected, the mutation N501Y, present in all three VOCs and also noted in other lineages such as A.27, makes its local environment in the RBD more neutral and hides negative charges exposed in the original (or B.1) RBD. This appeared further pronounced with the E484K change found in the RBD of both Beta and Gamma viruses. Indeed, this change is also located in the ACE2 binding interface, resulting in a more positively charged RBD surface, in turn contributing to better binding to the negatively charged patch in mACE2.

## DISCUSSION

The worldwide dissemination of SARS-CoV-2 with over 250 million confirmed cases, has been accompanied with the emergence of a multitude of variants bearing a constellation of changes in the ~30 kb viral genome. Some of these variants carry changes, often in the spike, which have been associated with increase in transmissibility or pathogenicity in humans (Barton et al., 2021; Harvey et al., 2021; Volz et al., 2021). Among those, the N501Y mutation has repeatedly appeared independently in the pandemic, and is notably shared by the three first VOCs. This change has been shown to increase the affinity of the spike protein for hACE2 (Starr et al., 2020; Tian et al., 2021; Zahradnik et al., 2021), which could increase transmissibility (Liu et al., 2021).

Our results, both *in vitro* and *in vivo*, demonstrate that SARS-CoV-2 variants carrying the N501Y mutation can efficiently replicate in mice. This host range expansion appears to be determined by N501Y, alone or in combination with additional mutations in the spike, such as those characteristic of the VOCs, but independently from the D614G mutation which is absent in the A.27 lineage. However, we noted that the viral titers of Alpha in the mice lungs were lower than those measured with other N501Y variants, suggesting that one or more mutations in the Alpha constellation negatively impact the virus fitness in mice. Interestingly, an Alpha isolate carrying the E484K substitution, present in Beta in Gamma, yielded lung viral titers comparable to other N501Y variants. This result is consistent with our structural modeling data indicating that E484K further increases the compatibility of the RBD with mACE2, and with recent results obtained with pseudotyped viruses showing that the N501Y, E484K and their combination increase entry in mACE2 expressing cells (Li et al., 2021).

Further in-depth studies will be needed to characterize the pathological consequences of infection with these variants. Notably, a mouse adapted strain carrying N501Y (Gu et al., 2020), in combination with two other substitutions in the spike (Sun et al., 2021) was reported to induce respiratory symptoms and lung inflammation with age-related mortality, while the young adult mice infected with our variants showed no signs of disease, except BALB/c mice infected with Beta which developed transient body weight loss. Thus, other changes among the constellations defining the VOC lineages might also play a role in the resulting *in vivo* phenotype. Indeed, most of the reported mouse-adapted variants also contain genetic changes outside of the spike. Further studies are needed to dissect the combinatory role of the mutations defining the SARS-CoV-2 VOCs, as well as their pathogenicity in older mice, or in other host genetic backgrounds. The ability of viruses of the N501Y-bearing lineages to replicate in common laboratory mice will facilitate *in vivo* studies in this species, to evaluate countermeasures (vaccines or therapeutic interventions), to assess antibody cross-reactivity and vaccine cross-protection (Martinez et al., 2021), and for functional studies using genetically altered mouse strains.

It has been proposed that persistence can select for host range expansion of animal viruses, by selecting for virus variants that recognize phylogenetic homologues of the receptor (Baric et al., 1999; Schickli et al., 1997). Interestingly, there is still speculation on the mechanism of emergence of the VOCs, which are all characterized by an unusually large number of mutations compared to their last common ancestor, including a number of changes (substitution and deletions) in the spike protein. While the genomic surveillance could have missed evolutionary intermediates leading to these lineages, similar patterns of accumulated changes were noted in some longitudinal studies of immunocompromised individuals infected by SARS-CoV-2 for extended periods of time (Choi et al., 2020; Kemp et al., 2021), leading to the hypothesis of a role of such long-term infections in their emergence. As mutations in positions of interest in the spike RBD (417, 484 and 501), alone or in combination, have been noted in other lineages during the extensive circulation of SARS-CoV-2 in human populations (2021), and associated *in vitro* with modification of the affinity for human ACE2, the observed host range expansion nevertheless likely represents only a serendipitous by-product of selection for increased transmissibility of SARS-CoV-2 in its current host.

To evaluate the consequence of host range expansion for wild mice, we assessed the possibility of transmission by close contact. We did not test indirect transmission by aerosols since mice, unlike ferrets or minks, do not efficiently transmit respiratory viruses by this route. We demonstrated that the Beta VOC could be transmitted not only from an infected mouse to a first-level contact individual, but also from first-level to second-level contact mice. Almost all published studies have considered only transmission from an experimentally infected individual to a first contact, which does not reflect the situation which may occur in wild rodent groups since the infectious dose used for intranasal inoculation is likely much higher that the dose an animal may be accidentally exposed to. One exception is a study on deer mice (*Peromyscus maniculatus*) which reported inconsistent virus detection in oral swabs after two passages (Griffin et al., 2021). Due to the low reliability of oral-pharyngeal swabs in mice for assessing the presence of the virus (which may reside mostly in the lower respiratory tract), we used antibody titers against the spike as an indicator of viral exposure. Although not proportional to viral load, antibody levels reflect the intensity of viral exposure. As expected, infected mice showed very high levels of anti-spike antibodies. The observation that Contact 2 mice had similar levels as Contact 1 mice suggests that they had been exposed to virus doses in the same range of magnitude. Although not yet established, it is likely that the infectious dose for mice of these new variants is low. Moreover, SARS-CoV-2 infection from exposed mice (Contact 1 to Contact 2) occurred in the absence of any clinical signs of illness, consistent with the observation of frequent inter-human transmission from asymptomatic infected individuals (Muller, 2021).

Our results clearly demonstrate that, once they have been exposed to the virus, mice can transmit the virus to others, which raises major questions on the risk of mice or other rodents living in very large numbers in proximity to humans, of becoming secondary reservoirs for SARS-CoV-2 in regions where the N501Y-carrying variants circulate, from where they could evolve separately and potentially spillback to humans. Indeed, rodents have been hypothesized as the ancestral host of some betacoronaviruses (lineage A, which includes the seasonal human coronaviruses OC43 and HKU1 (Lau et al., 2015; Tsoleridis et al., 2019)). Similar and actionable concerns were raised upon the detection of SARS-CoV-2 in Mink farms in The Netherlands (Oreshkova et al., 2020) and in Denmark (Lassaunière R et al., 2020) due to the density of animals housed, and the detection of changes in the virus genome. While rodent densities are highly variable and more difficult to estimate, multiple spillovers have been detected in free-living White-tailed deer populations in North America (Kuchipudi et al., 2021). We posit that host range should be closely monitored along the continued evolution of SARS-CoV-2.

## Acknowledgments

We are grateful to Dr Luis Enjuanes (National Center for Biotechnology, Spain) for the generous gift of the DBT cells expressing mACE2. We thank Hélène Huet and Jean-Luc Servely (Unité d’Histologie et d’Anatomie pathologique, Ecole nationale vétérinaire d’Alfort) for the histological and immunohistochemical techniques, and the histology pictures, respectively. We also thank the team of the core facility P2M (Institut Pasteur) for genomic sequencing. We acknowledge the authors, originating and submitting laboratories of the sequences from GISAID (Suppl. Table 1). We avoided any direct analysis of genomic data not submitted as part of this paper and used this genomic data only as background. This work used the computational and storage services (Maestro cluster) provided by the IT department at Institut Pasteur, Paris. FG is part of the Pasteur-Paris University (PPU) International PhD programme, BioSPC doctoral school.

This work was supported by the « URGENCE COVID-19 » fundraising campaign of Institut Pasteur), the French Government’s Investissement d’Avenir program, Laboratoire d’Excellence Integrative Biology of Emerging Infectious Diseases (Grant No. ANR-10-LABX-62-IBEID), the Agence Nationale de la Recherche (Grant No. ANR-20-COVI-0028-01) and the RECOVER project funded by the European Union’s Horizon 2020 research and innovation programme under grant agreement No. 101003589. ESL acknowledges funding from the INCEPTION programme (Investissements d’Avenir grant ANR-16-CONV-0005).

## Author contributions

XM and ESL designed and coordinated the study. MP, LL and LC performed in vitro experiments and viral quantification. XM, LC and JJ performed in vivo experiments. GJ performed histopathological analysis. EBS and FAR performed the structural analysis. ESL and FG performed the phylogenetic analysis. FD and MA isolated and produced the viral isolates under the supervision of SB, VE and SvdW. XM and ESL wrote and revised the manuscript with input from all authors.

## Materials and Methods

### Cells and viruses

VeroE6 cells (African green monkey kidney cells from ATCC (CCL-81)), A549-hACE2 cells (human adenocarcinoma alveolar epithelial cells expressing the human ACE2 receptor, a kind gift of Olivier Schwartz) and DBT-mACE2 cells (a kind gift of Dr Luis Enjuanes) were maintained in Dulbecco’s Modified Eagle’s Medium (DMEM) supplemented with 10% fetal bovine serum (FBS) and penicillin/streptomycin. A549-hACE2 cells were supplemented with 10 μg/ml of Blasticidin and DBT-mACE2 cells with 800 μg/ml of G418 to maintain the plasmid expressing ACE2.

All SARS-CoV-2 isolates were supplied by the National Reference Centre for Respiratory Viruses hosted by Institut Pasteur (Paris, France) and headed by Pr. S. van der Werf. The human sample from which hCoV-19/France/GES-1973/2020 was isolated was provided by Pr. Laurent Andreoletti, CHU of Reims, France. As previously described (Planas et al., 2021), the human sample from which strain hCoV-19/France/IDF-IPP158i/2020 was isolated has been provided by Dr. Karl Stefic et Pr. Catherine Gaudy Graffin, CHRU de Tours, Tours, France and the human sample from which strain hcoV-19/France/IDF-IPP00078/2021 was isolated has been provided by Dr. Mounira Smati-Lafarge, CHI of Créteil, Créteil, France. The human sample from which hCoV-19/France/IDF-IPP05174/2021 was isolated was provided by Dr. Marianne Leruez-Ville, Necker hospital, Paris, France. The human sample from which hCoV-19/France/HDF-IPP11602/2021 was isolated was provided by Dr. Raphaël Guiheneuf, CH Simone Veil, Beauvais, France. The human sample from which strain hCoV-19/FrenchGuiana/IPP03772/2021 was isolated has been provided by Dr. Dominique Rousset, Institut Pasteur de la Guyane. This P.1 lineage virus was isolated by inoculation of VeroE6 cells, followed by two passages. Viruses were amplified and titrated by standard plaque forming assay on VeroE6 cells. The sequence of the stocks was verified by RNAseq on the mutualized platform for microbiology (P2M). All work with infectious virus was performed in biosafety level 3 containment laboratories at Institut Pasteur.

### Phylogenetic analysis

We used the Nextstrain (Hadfield et al., 2018) pipeline (https://github.com/nextstrain/ncov) to reconstruct a global, representatively subsampled phylogeny highlighting the position of the isolates used in the experimental work. A maximum likelihood phylogenetic tree was built based on the GTR model, after masking 130 and 50 nucleotides from the 5’ and 3’ ends of the alignment, respectively, as well as single nucleotides at positions 18529, 29849, 29851, 29853. We checked for temporal signal using Tempest v1.5.3 (Rambaut et al., 2016). The temporal phylogenetic analyses were performed with augur and TreeTime (Sagulenko et al., 2018), assuming a clock rate of 830.0008±0.0004 (SD) substitutions/site/year (Rambaut, 2020), coalescent skyline population growth model and the root set on the branch leading to the Wuhan/Hu-1/2020 sequence. The time and divergence trees were visualized with FigTree v1.4.4 86 (http://tree.bio.ed.ac.uk/software/figtree/).

### *In vivo* studies

Eight-week-old female BALB/cJRj (BALB/c) and C57BL/6JRj (C57BL/6) mice were purchased from Janvier Labs (Le Genest St Isle, France). Infection studies were performed in animal biosafety level 3 (BSL-3) facilities at the Institut Pasteur, in Paris. All animal work was approved by the Institut Pasteur Ethics Committee (projects dap200008 and dap21050) and authorized by the French Ministry of Research under projects 24613 and 31816 in compliance with the European and French regulations on the protection of live animals. Throughout experiments, all mice were maintained in a 22-24°C, 14:10hr day-night environment, and received food and water ad libitum Anesthetized (ketamine/xylazine) mice were inoculated intranasally with 6 x10^4^ PFU of SARS-CoV-2 variants in 40-44μl volume. Clinical signs of disease and weight loss were monitored daily. Mice were euthanized by ketamine/xylazine overdose at indicated time points (–dpi 2, 3 or 4) when samples for titer and histopathological analyses were collected. The left lung lobe was fixed by submersion in 10% phosphate buffered formalin for 7 days prior to removal from the BSL3 for processing and transfer in 70% ethanol. The right lung lobe was placed on a 70μ cell strainer (Falcon), minced with fine scissors and ground with a syringe plunger using 400 μl of PBS. Lung homogenates were used for viral quantification.

To assess transmission, two 15-week-old BALB/c female mice were inoculated intranasally with 6×10^4^ PFU of Beta SARS-CoV-2 variant (see Fig. 2). On the following day (dpi1), three naive mice (Contact 1, 15-week-old BALB/c females) were introduced in their cage. One Contact 1 mouse was removed on dpi3 and placed in a cage containing three other naive mice (Contact 2, 15-week-old BALB/c females). The last two Contact 1 mice were separated from the infected mice on dpi4 and put in cage 2 with the other Contact 1 and the Contact 2 mice. Fourteen days later (dpi18), all mice were bled to assess their anti-S antibody levels by ELISA. The experiment was performed on three such cohorts.

### Anti-S ELISA

High-binding 96-well ELISA plates (Costar, Corning) were coated overnight with purified SARS-CoV-2 S protein in the form of a trimer (125 ng/well in PBS). After washings with 0.1% Tween 20 in PBS (PBST), plates were blocked for 2h with PBS added with 1% Tween 20, 5% sucrose and 3% milk powder (Blocking solution). After PBST washings, 1:100-diluted mouse sera (in PBST + 1% BSA) and 7 consecutive 1:3 dilutions were added and incubated for 2 h. After washings, plates were revealed by addition of goat HRP-conjugated anti-mouse IgG (0.8 μg/ml final in blocking solution, Immunology Jackson ImmunoReseach). Plates were revealed by adding 100 μl of HRP chromogenic substrate (ABTS solution, Euromedex) after PBST washings. Experiments were performed in duplicate at room temperature using a HydroSpeed microplate washer and Sunrise microplate absorbance reader (Tecan, Mannedorf), with OD at 405nm. Area under the curve (AUC) values were determined by plotting the log10 of the dilution factor values (X axis) required to obtain OD405nm values (Y axis). AUC calculation analyses were performed using GraphPad Prism software (v8.4.1, GraphPad Prism Inc.).

### Histopathological analysis

Histological analysis was performed on paraffin-embedded 4μm-thick sections used for hematoxylin-eosin (H&E) staining and for the immunohistochemical detection of the virus, using a rabbit polyclonal primary antibody directed against SARS-CoV nucleocapsid (N) protein (Novus Biologicals, cat #NB100-56576; dilution: 1:200). IHC was carried out based on the manufacturer’s recommendations.

### Virus quantification

Tissue homogenates were aliquoted for RNA quantification and titration. Viral RNA was extracted using the QIAamp viral RNA mini kit (Qiagen). Viral RNA quantification was performed by quantitative reverse transcription PCR (RT-qPCR) using the IP4 set of primers and probe as described on the WHO website (https://www.who.int/docs/default-34source/coronaviruse/real-time-rt-pcr-assays-for-the-detection-of-sars-cov-2-institut-35pasteur-paris.pdf?sfvrsn=3662fcb6_2) and the Luna Universal Probe One-Step RT-qPCR Kit (New England Biolabs, France).

For plaque assay, 10-fold serial dilutions of samples in DMEM were added onto VeroE6 monolayers in 24 well plates. After one-hour incubation at 37°C, the inoculum was replaced with 2% FBS DMEM and 1% Carboxymethylcellulose. Three days later, cells were fixed with 4% formaldehyde, followed by staining with 1% crystal violet to visualize the plaques.

### Structural analysis

In order to study the effect on mACE2 binding of mutations in the RBD of the spike protein of the new SARS-CoV-2 variants, we generated a homology model for mACE2/RBD complex based on the available X-ray structure of the hACE2/RBD complex and analyzed their electrostatic surface potential at neutral pH. We used this model to analyze the predicted electrostatic potential at the ACE2/RBD interface. We generated homology models of mACE2/RBD complexes corresponding to the various SARS-CoV-2 variants using the MODELLER software (Webb and Sali, 2016) based on the crystal structure of hACE2 peptidase domain in complex with SARS-CoV-2 spike RBD (PBD 6M0J). The quality of the models was inspected using PROCHECK and MOLPROBITY servers (Laskowski et al., 2012; Williams et al., 2018). The surface electrostatic potential of the various models was calculated using the APBS/PDB2PQR server at pH 7.0 using a PARSE forcefield. The cartoon diagrams and surface representations were generated using PyMOL (Schrödinger, LLC)

### Statistical analysis

Statistical analysis and graphing of data were performed using R software. Viral loads and titers and serology data were first analyzed by one-way ANOVA and groups were then compared with Tukey’s test.

## Supplemental information

**Fig. S1.**
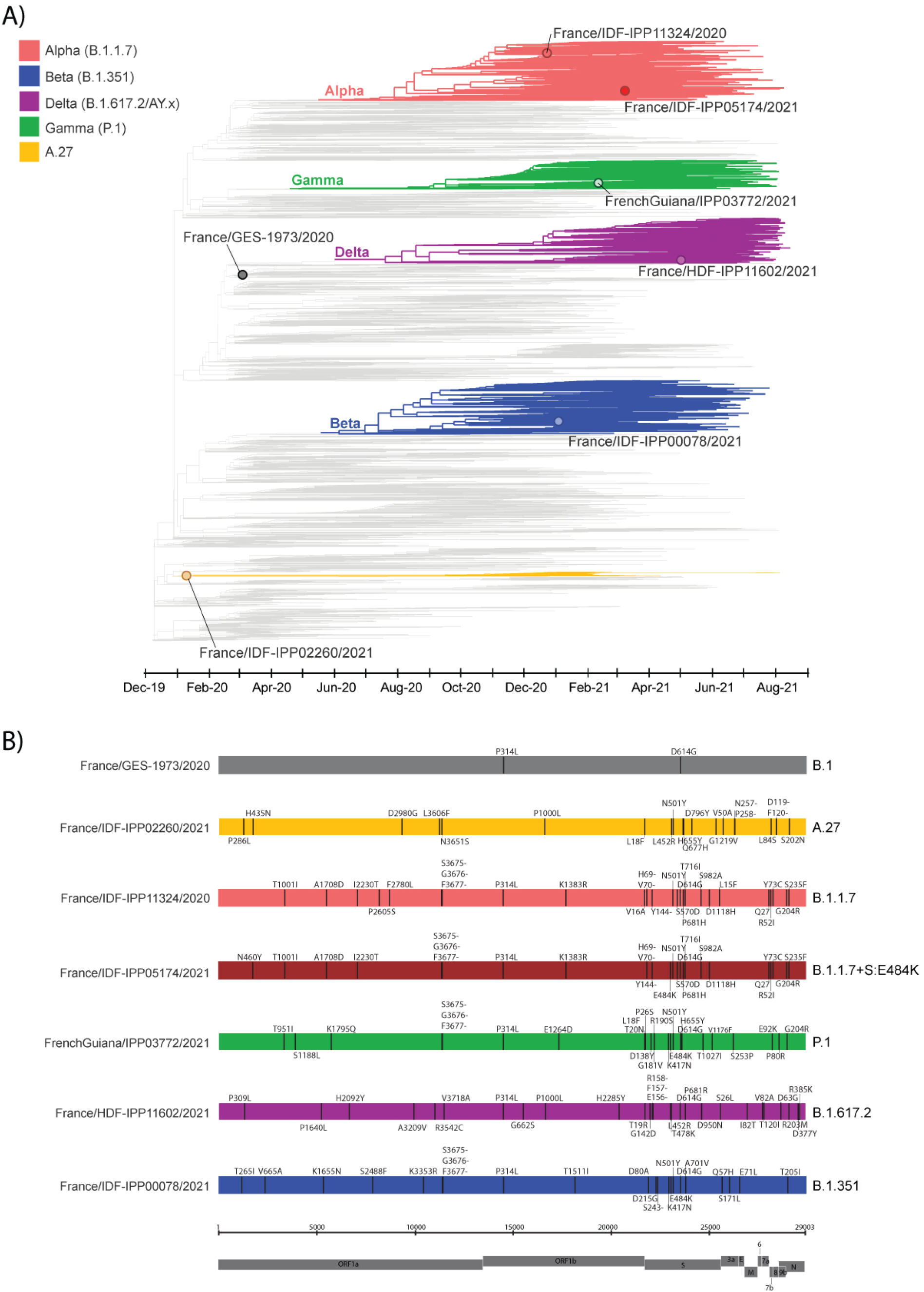
SARS-CoV-2 isolates in a global context. Maximum likelihood phylogeny where the sequence corresponding to each variant is highlighted. Scheme of the variants in comparison to the ancestral sequence (NC_045512).

**Fig. S2.**
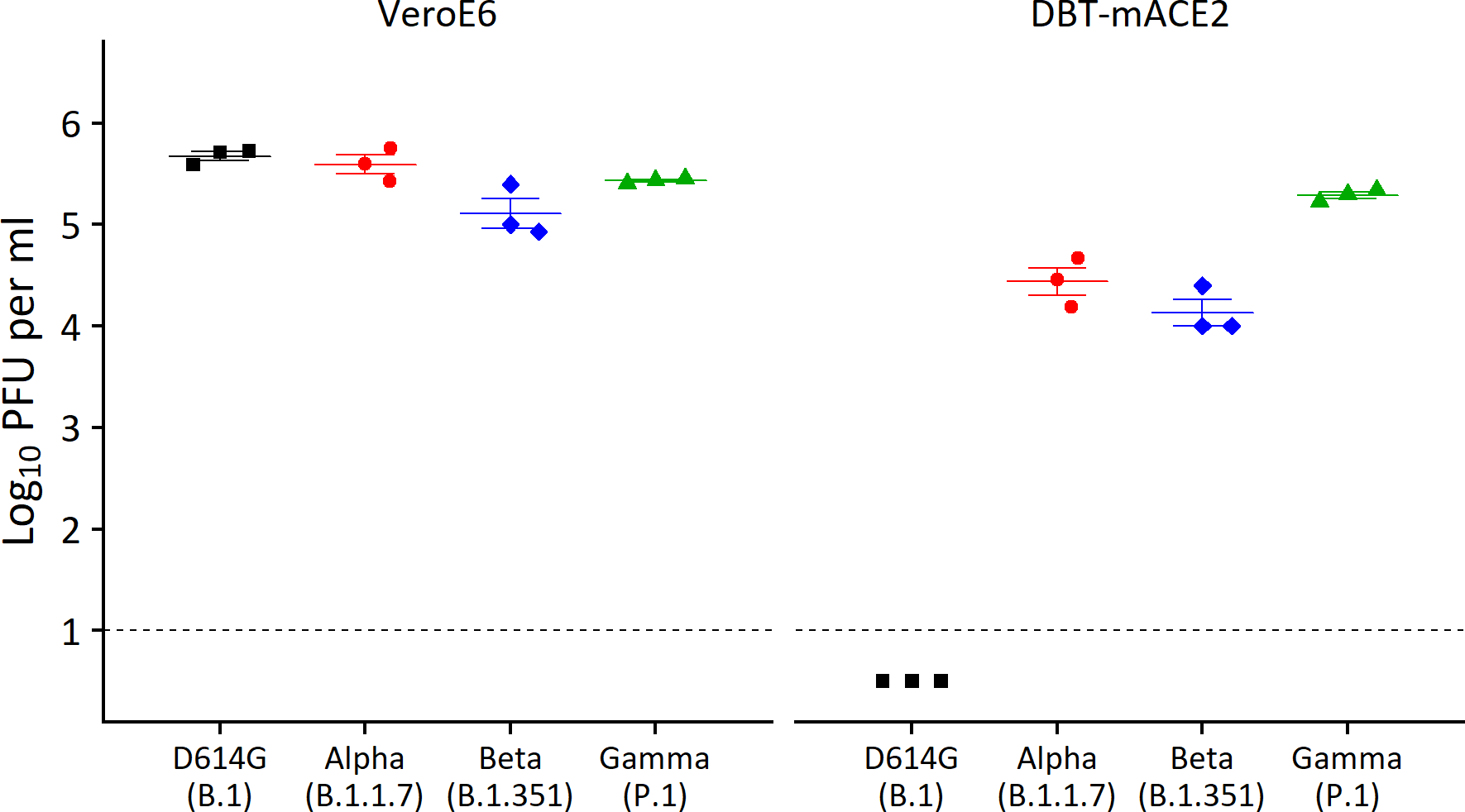
SARS-CoV-2 N501Y variants show strong replication in VeroE6 cells and in a mouse cell line expressing mACE2. Viral titer in cell supernatant 48h after infection of Vero-E6 cell and DBT-mACE2 cell lines at a MOI = 0.1. Viruses were titrated on VeroE6 cells. The dotted line represents LOD, and undetected samples are plotted at half the LOD. While the B.1 virus did not replicate in DBT-mACE2 cells, all variants reached very high titers.

**Fig. S3.**
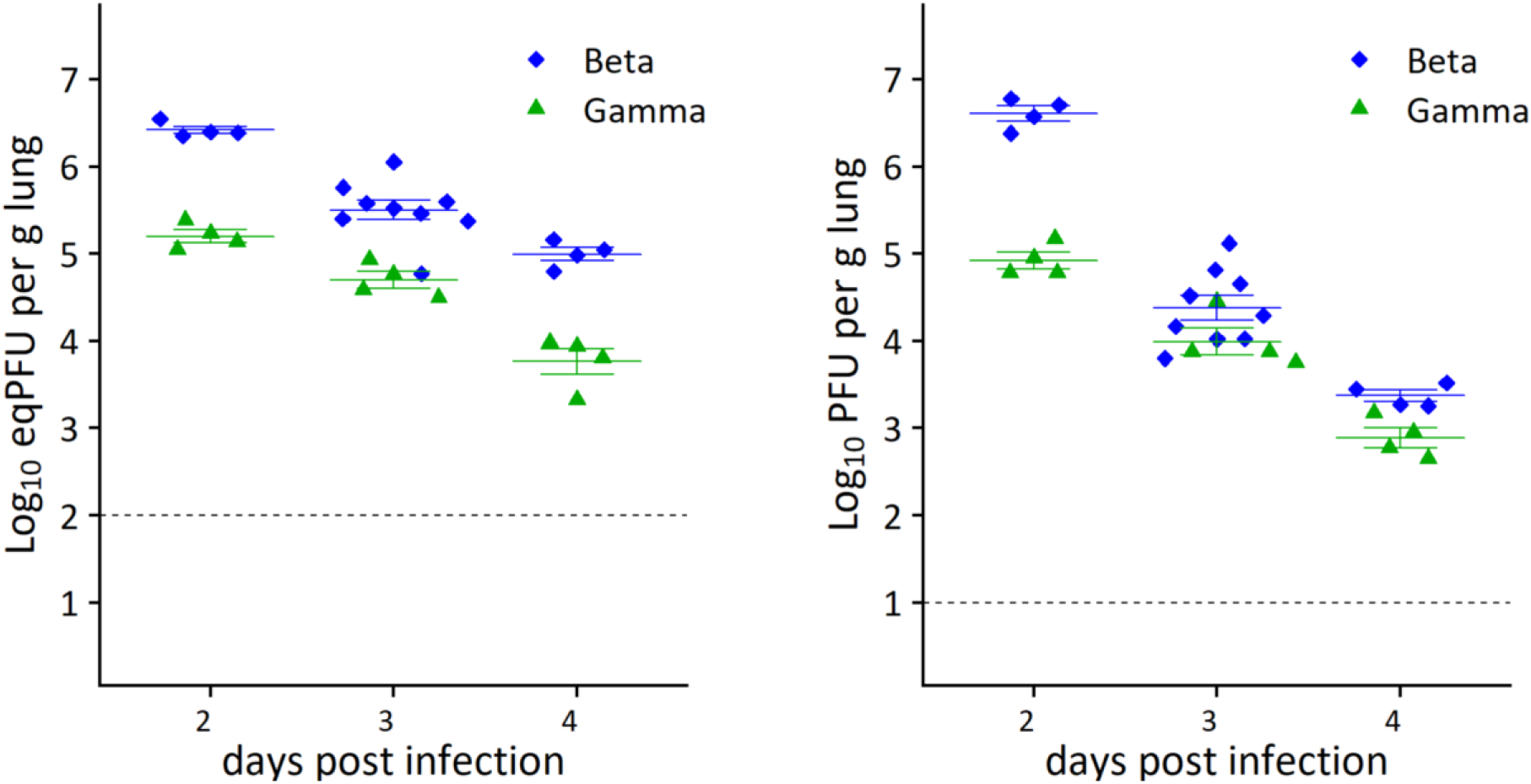
Replication of SARS-CoV-2 B.1.351 and P.1 variants in lungs peaks at day 2 post inoculation in young adult BALB/c mice. Mice were inoculated intranasally with 6×10^4^ PFU of SARS-CoV-2 variants. Lungs were harvested at dpi 2, 3 and 4. Viral load was quantified by RT-qPCR (left). Viral titer was quantified on VeroE6 cells (right). The dotted line represents LOD.

**Fig. S4.**
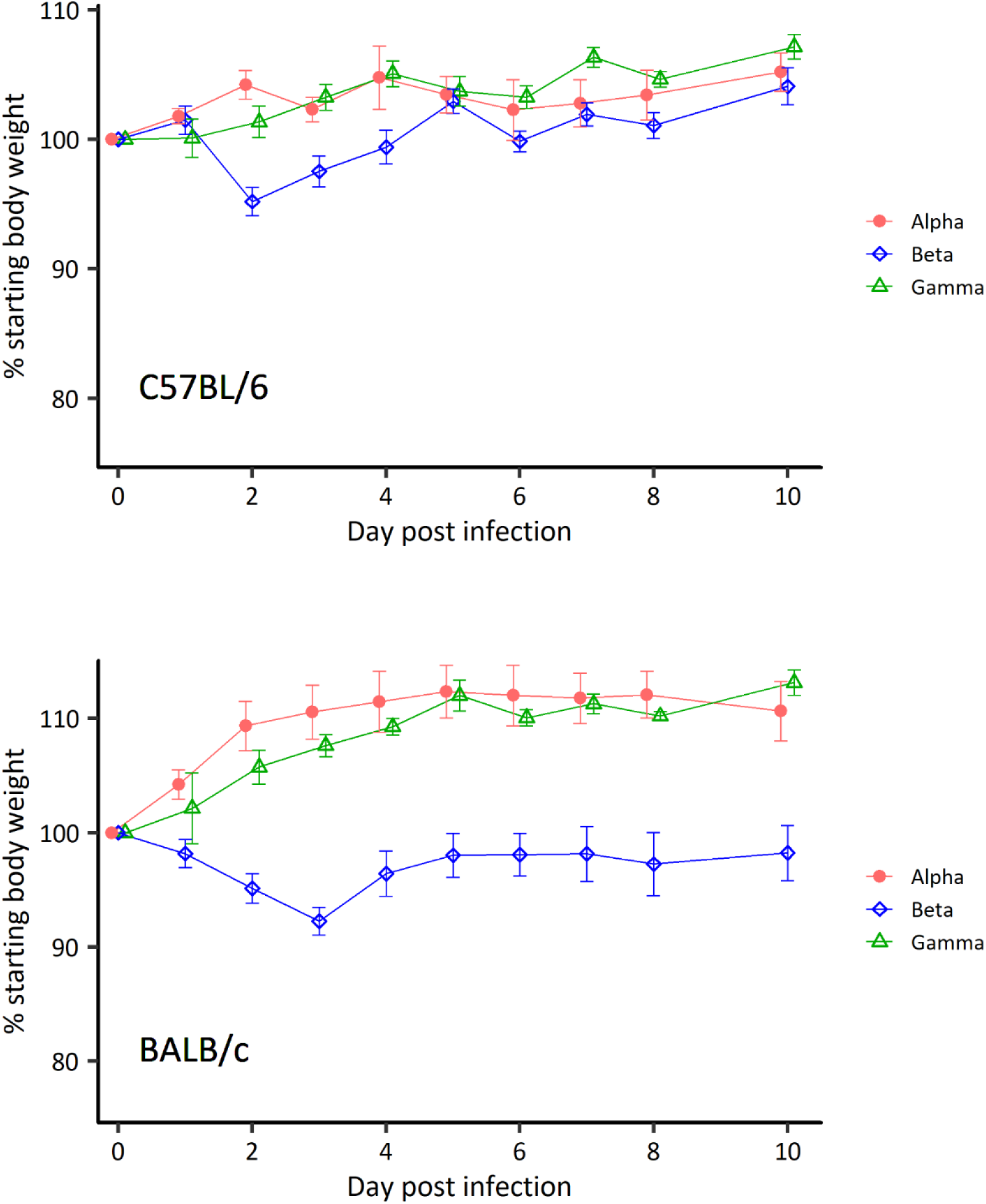
Body weight variations in BALB/c and C57BL/6 mice inoculated with SARS-CoV-2 alpha, beta and gamma variants. Mice were inoculated intranasally with 6×10^4^ PFU of SARS-CoV-2 variants. Mild to moderate and transient body weight loss was observed with a peak on dpi 2 or 3 only with the beta variant.

**Suppl Table 1.** GISAID acknowledgements.

**Suppl Table 2.** Viruses.

| Pango lineage | Strain | GISAID accession Id |
| --- | --- | --- |
| B.1 | hCoV-19/France/GES-1973/2020 | EPI_ISL_414631 |
| B.1.1.7 | hCoV-19/France/IPP-00158i/2021 | EPI_ISL_1259025 |
| B.1.1.7+S:E484K | hCoV-19/France/IDF-IPP05174/2021 | EPI_ISL_1381817 |
| B.1.351 | hCoV-19/France/IDF-IPP00078/2021 | EPI_ISL_964916 |
| P.1 | hCoV-19/FrenchGuiana/IPP03772/2021 | EPI_ISL_1096262 |
| B.1.617.2 | hCoV-19/France/HDF-IPP11602/2021 | EPI_ISL_2188419 |
| A.27 | hCoV-19/France/IDF-IPP02260/2021 | EPI_ISL_4211287 |

